# C/EBPβ regulates hypertrophic versus hyperplastic fat tissue growth

**DOI:** 10.1101/2020.09.02.278911

**Authors:** Christine Müller, Laura M. Zidek, Sabrina Eichwald, Cornelis F. Calkhoven

## Abstract

Chronic obesity is correlated with severe metabolic and cardiovascular diseases as well as with an increased risk for developing cancers. Obesity is usually characterized by fat accumulation in enlarged - hypertrophic – adipocytes that are a source of inflammatory mediators, which is seen as causal for developing metabolic disorders. Yet, in certain healthy obese individuals fat is stored in metabolically more favorable *hyperplastic* fat tissue that contains an increased number of smaller adipocytes that are less inflamed. In a previous study we demonstrated that C/EBPβ-LIP deficient, yet C/EBPβ-LAP proficient mice show an expanded health and lifespan. Here we show that in mice on a high-fat diet LIP-deficiency results in adipocyte hyperplasia as well as efficient fat storage in subcutaneous depots associated with metabolic and inflammatory improvements. Our data identify C/EBPβ as a regulator of adipocyte fate in response to increased fat intake, which has major implications for metabolic health and aging.

## Introduction

Nutrient overload, particularly in combination with a sedentary lifestyle, is the main cause of the increasing incidence of obesity we are facing today. In most cases chronic obesity provokes the development of metabolic diseases like insulin resistance, type 2 diabetes (T2D), non-alcoholic fatty liver disease (NFALD) and non-alcoholic steatohepatitis (NASH) (Longo et al., 2019). The surplus of fat is stored in the white adipose tissue (WAT) largely without an increase in adipocyte number, resulting in an increase in fat cell size. This hypertrophy is accompanied with reduced vascularization and oxygen supply, and an increase in macrophage infiltration, and inflammation (Frasca et al., 2017; Tchkonia et al., 2010). Since the storage capacity of the hypertrophic cells is limited, fat starts to accumulates in ectopic tissues like liver, heart and skeletal muscle (Frasca et al., 2017). This dyslipidemia, also referred to as “lipotoxicity” further promotes metabolic disorders. However, there is an exception from this scenario as individuals exist that are chronically obese but stay - at least transiently - metabolically healthy. Evidence from mouse and a few human studies suggest that storing surplus of nutrients as fat through adipocyte hyperplasia (increasing number) is associated with metabolic health (White and Ravussin, 2019). As the fat storage can be distributed over more fat cells, the individual fat cells stay smaller, are metabolically more active, and less inflamed (Ghaben and Scherer, 2019). So far not much is known about the underlying molecular mechanisms and involved regulators that drive fat storage in either the hypertrophic or the hyperplastic direction. Such regulators may be attractive targets to therapeutically switch the adipocytes from a hypertrophic into a hyperplasic state in order to prevent metabolic complications associated with obesity. CCAAT/Enhancer Binding Protein beta (C/EBPβ) is a transcription factor known to regulate adipocyte differentiation together with other C/EBPs and peroxisome proliferator-activated receptor gamma (PPARγ) (Siersbaek and Mandrup, 2011). In all cases of C/EBPβ controlled cellular processes it is important to consider that different protein isoforms of C/EBPβ exist. The two long C/EBPβ isoforms, LAP1 and LAP2 (Liver-enriched activating proteins) differ slightly in length and both function as transcriptional activators. The N-terminally truncated isoform LIP (Liver-enriched inhibitory protein) acts inhibitory because it lacks transactivation domains yet binds to DNA in competition with LAP1/2 (Descombes and Schibler, 1991). We have shown earlier that LIP expression is stimulated by mTORC1 signaling involving a *cis*-regulatory short upstream open reading frame (uORF) in the Cebpb-mRNA (Calkhoven et al., 2000; Zidek et al., 2015).

Mutation of the uORF in mice (*Cebpb*^*ΔuORF*^ mice) results in loss of LIP expression, unleashing LAP transactivation function, resulting in C/EBPβ super-function. (Müller et al., 2018; Wethmar et al., 2010; Zidek et al., 2015) In *Cebpb*^*ΔuORF*^ mice metabolic and physical health is preserved and maintained during ageing with features also observed under calorie restriction (CR), including leanness, enhanced fatty acid oxidation, prevention of steatosis, better insulin sensitivity and glucose tolerance, preservation of motor coordination, delayed immunological ageing and reduced interindividual variation in gene expression of particularly metabolic genes (Müller et al., 2018; Zidek et al., 2015).

Here we show that C/EBPβ is critically involved in dictating the adipocyte phenotype and the metabolic outcome in response to high-fat diet (HFD) feeding in mice. The *Cebpb*^*ΔuORF*^ mice accumulate fat in hyperplastic rather than hypertrophic depots. In addition, *Cebpb*^*ΔuORF*^ mice store the surplus of fat more efficiently in subcutaneous fat stores. Consequently, the *Cebpb*^*ΔuORF*^ mice better maintain glucose tolerance and they are protected against the development of dyslipidemia, indicating a healthy obese phenotype.

## Results

*Cebpb*^*ΔuORF*^ and wild-type (wt) control littermates on a C57BL/6J background were fed a high-fat diet (HFD; 60% fat) or standard chow (10% fat) for a period of 19 weeks. Both genotypes gained weight over the whole experimental period with significantly lower body weights for *Cebpb*^*ΔuORF*^ mice, suggesting that they are partially protected from HFD induced weight gain (**Figure 1A**). Both the food intake and the energy efficiency (energy that was extracted from the food during digestion) were similar in *Cebpb*^*ΔuORF*^ and wt mice (**Figure 1 – figure supplement 1A, B**). Lean mass and fat mass body composition was determined by computer tomography (CT) after 19 weeks of HFD feeding. On standard chow, fat mass of *Cebpb*^*ΔuORF*^ mice is reduced compared to wt mice (**Figure 1B**) as we showed before (Zidek et al., 2015). Unexpectedly, we observed a slight but significantly increased overall fat mass in the *Cebpb*^*ΔuORF*^ mice compared to wt controls (**Figure 1B**), which correlates with a relative reduction in lean mass (**Figure 1 – figure supplement 1C**). There was no significant difference in the amount of visceral fat mass between wt and *Cebpb*^*ΔuORF*^ mice under HFD (**Figure 1C**). However, the HFD fed *Cebpb*^*ΔuORF*^ mice had a significantly increased subcutaneous fat mass compared to the wt controls (**Figure 1D**). Take together the data show that *Cebpb*^*ΔuORF*^ mice store more fat upon HFD feeding compared to wt littermates, but this extra fat is mainly stored in the subcutaneous fat depot.

**Figure 1.**
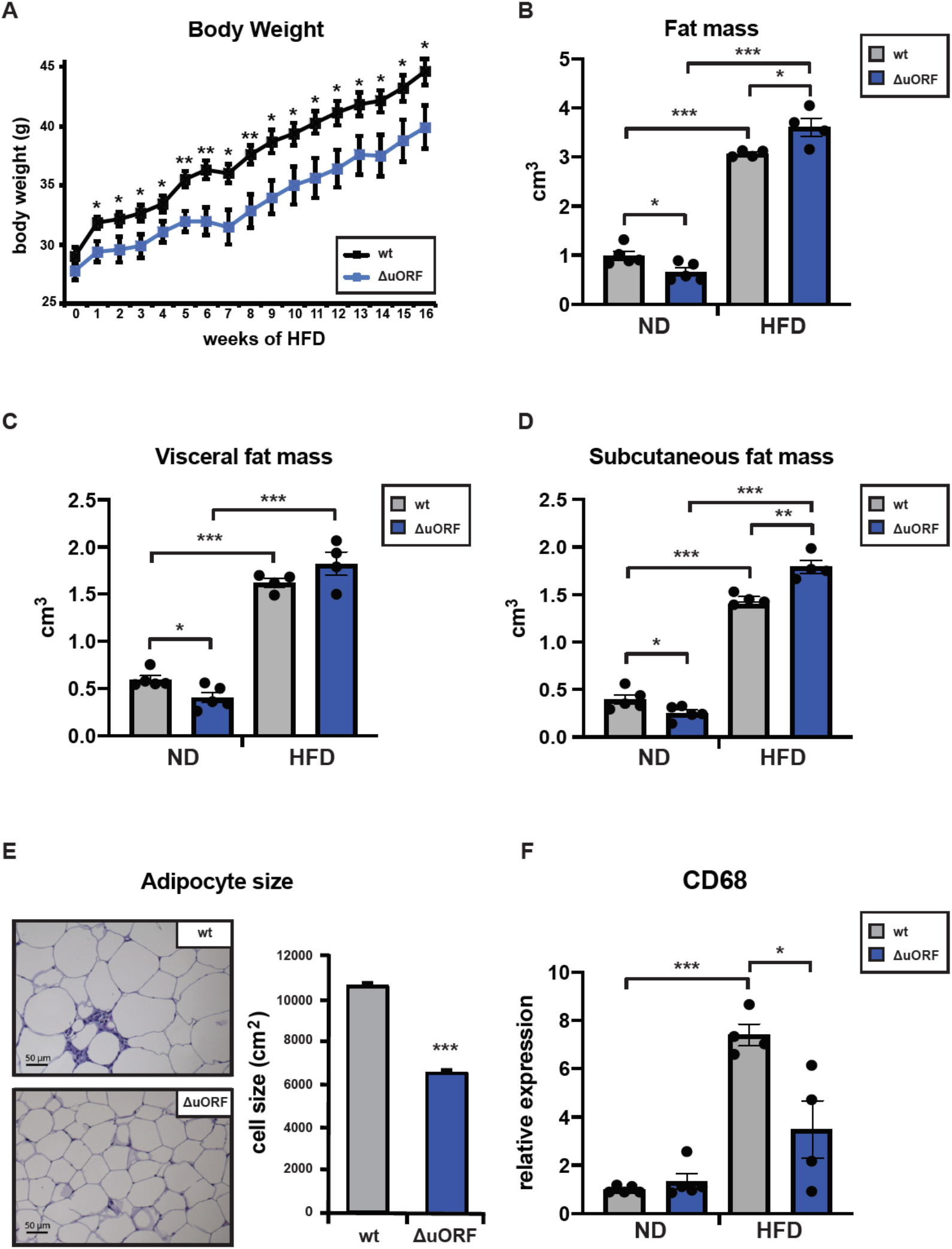
*Cebpb*^*ΔuORF*^ mice on high-fat diet (HFD) store fat in hyperplastic adipocytes and the subcutaneous depot. (A) Growth curves of wt and *Cebpb*^*ΔuORF*^ mice (ΔuORF) on HFD (wt, n = 10; *Cebpb*^*ΔuORF*^, n = 8). (B) Volume of total fat mass as measured by abdominal CT analyses (19 weeks; wt, n = 5; ΔuORF n = 4). (C) Volume of visceral fat mass as measured by abdominal CT analyses (19 weeks; ND, n = 5; HFD, n = 4) (D) Volume of subcutaneous fat mass as measured by abdominal CT analyses (19 weeks; ND, n = 5; HFD, n = 4) (E) Histological hematoxylin and eosin (H&E) staining of epididymal WAT (19 weeks; scale bar corresponds to 50 μm). Quantification of the fat cell area is shown at the right (n = 6, 10 adjacent cells are measured per mouse). (F) Relative expression of CD68 macrophage marker as determined by qPCR (19 weeks HF; wt, n = 5; ΔuORF n = 4). All values are mean ± SEM. P-values were determined with Student’s t-test, *P < 0.05; **P < 0.01; ***P < 0.001.

Next we compared histological sections from visceral fat of HFD fed *Cebpb*^*ΔuORF*^ mice and wt littermates and observed that the adipocyte size in the *Cebpb*^*ΔuORF*^ mice is significantly smaller compared to the wt mice (**Figure 1E**). Since the absolute volume of the visceral fat is not different between the two genotypes the smaller adipocyte volume for *Cebpb*^*ΔuORF*^ mice means that these mice have a higher number of adipocytes in their visceral fat compared to wt mice. Calculated from fat volume (CT analysis) and cell size (histology) the *Cebpb*^*ΔuORF*^ mice have 1.84 times more fat cells in the visceral fat compartment than wt mice on HFD, and compared to ND fed mice *Cebpb*^*ΔuORF*^ mice increased their fat cell number on HFD by 2.37, while wt mice increased fat cell number only by 1.36 times (ND data were taken from (Zidek et al., 2015). Adipose tissue composed of small adipocytes is metabolically more active and better supplied with oxygen, and its inflammatory state is usually lower than that of enlarged adipocytes (Ghaben and Scherer, 2019). Analysis of the inflammatory state of visceral fat by determining the expression of the macrophage marker CD68 using quantitative PCR (qPCR) showed a significantly reduction in CD68 expression for *Cebpb*^*ΔuORF*^ mice compared to wt controls on HFD (**Figure 1F**). These data demonstrate that *Cebpb*^*ΔuORF*^ mice store extra fat through an increase in adipocyte numbers (hyperplasia), which results in smaller and healthier adipocytes and reduced fat tissue inflammation.

**Figure supplement 1.**
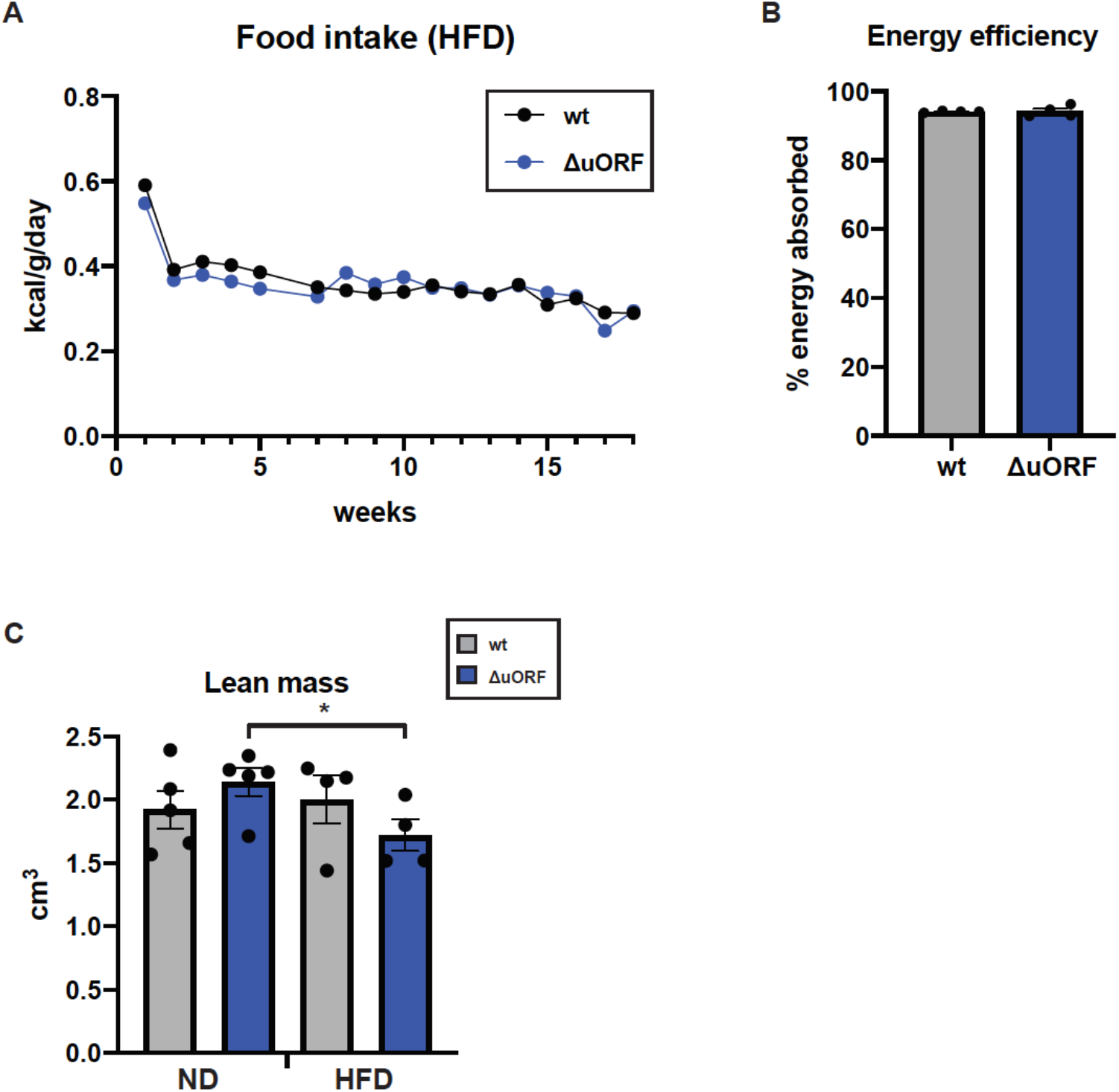
(A) Daily food intake per mouse, normalized to body weight as determined over 18 weeks (wt, n = 9 mice / 4 cages; ΔuORF n = 7 mice / 2 cages). (B) Efficiency of caloric utilization measured by bomb calorimetry of food and faces (wt, n = 4; ΔuORF n = 4). (C) Volume of lean body mass of *Cebpb*^*ΔuORF*^ mice measured by abdominal CT analyses (19 weeks; ND, n = 5; HFD n = 4). All values are mean ± SEM. P-values were determined with Student’s t-test, *P <0.05; **P < 0.01; ***P < 0.001.

Adipocyte hypertrophy under obese conditions is associated with a limit in fat storage capacity of the adipocytes and basal enhanced lipolysis (Khan et al., 2009; Laurencikiene et al., 2011) The resulting increase in the concentration of fatty acids in the circulation causes lipid accumulation in non-fat tissues like liver, muscle and heart (Longo et al., 2019). We stained histological sections of liver, muscle and heart with the lipid stain Sudan III to address differences in dyslipidemias. Compared to wt mice on a HFD the lipid accumulation is strongly reduced in *Cebpb*^*ΔuORF*^ mice, particularly in the liver and to a lesser extend in muscle and heart tissue. In agreement with the reduced dyslipidemia in *Cebpb*^*ΔuORF*^ mice on HFD the weight of their liver or heart is lower (**Figure 2 – figure supplement 1**). These data demonstrate that *Cebpb*^*ΔuORF*^ mice are protected against dyslipidemias in response to HFD.

**Figure 2.**
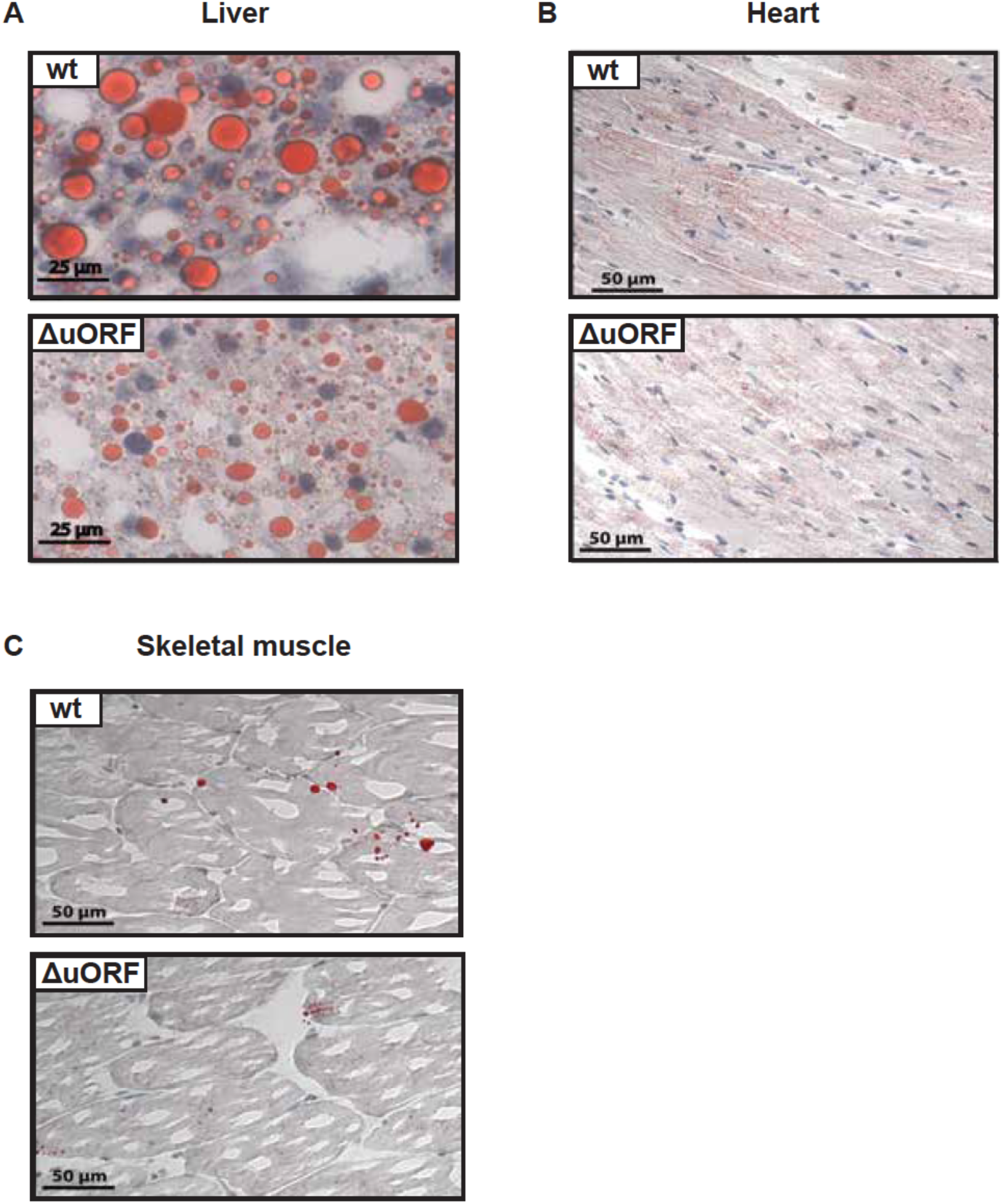
*Cebpb*^*ΔuORF*^ mice on high-fat diet (HFD) are protected against dyslipidemia. Histological sections of (A) liver, (B) cardiac muscle and (C) skeletal muscle of wt or *Cebpb*^*ΔuORF*^ mice (ΔuORF) (19 weeks). Sections were stained with hematoxylin (blue) and Sudan III for red color lipid staining.

**Figure supplement 1.**
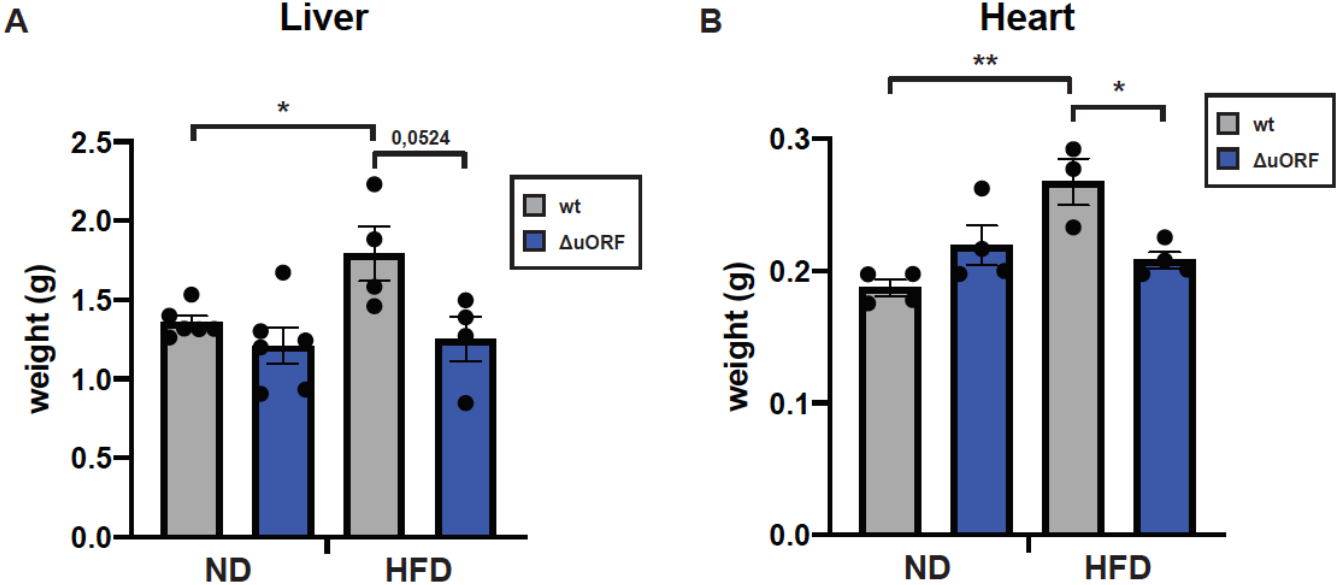
Liver and heart weights of *Cebpb*^*ΔuORF*^ mice and wt mice on normal diet (ND) or HFD (19 weeks) (Liver ND n=6; Liver HFD n=4; Heart HFD wt n=3, ΔuORF n=4). All values are mean ± SEM. P-values were determined with Student’s t-test, *P < 0.05.

Chronic obesity often results in the loss of glucose homeostasis (Abranches et al., 2015). We therefore analyzed glucose tolerance and insulin sensitivity in the HFD fed *Cebpb*^*ΔuORF*^ mice and wt littermates. Glucose clearance from the circulation measured by intraperitoneal glucose tolerance test (IPGTT) was impaired in response to 7 weeks HFD feeding for the wt mice, as is shown by a significantly increased area under the curve (AUC) (**Figure 3A**). For the *Cebpb*^*ΔuORF*^ mice the already significantly better glucose clearance on normal diet does not change on HFD. At this time point the intraperitoneal insulin sensitivity test (IPIST) did not show a difference between ND and HFD, apart from that the *Cebpb*^*ΔuORF*^ mice do perform better in general (**Figure 3B**). In conclusion, our data show that *Cebpb*^*ΔuORF*^ mice on HFD feeding perform better on glucose tolerance, are protected against lipodystrophy and show a lower inflammatory status of WAT. These metabolically favorable phenotypes of the *Cebpb*^*ΔuORF*^ mice correlate with hyperplastic fat storage as well as more efficient fat accumulation in the subcutaneous depot.

**Figure 3.**
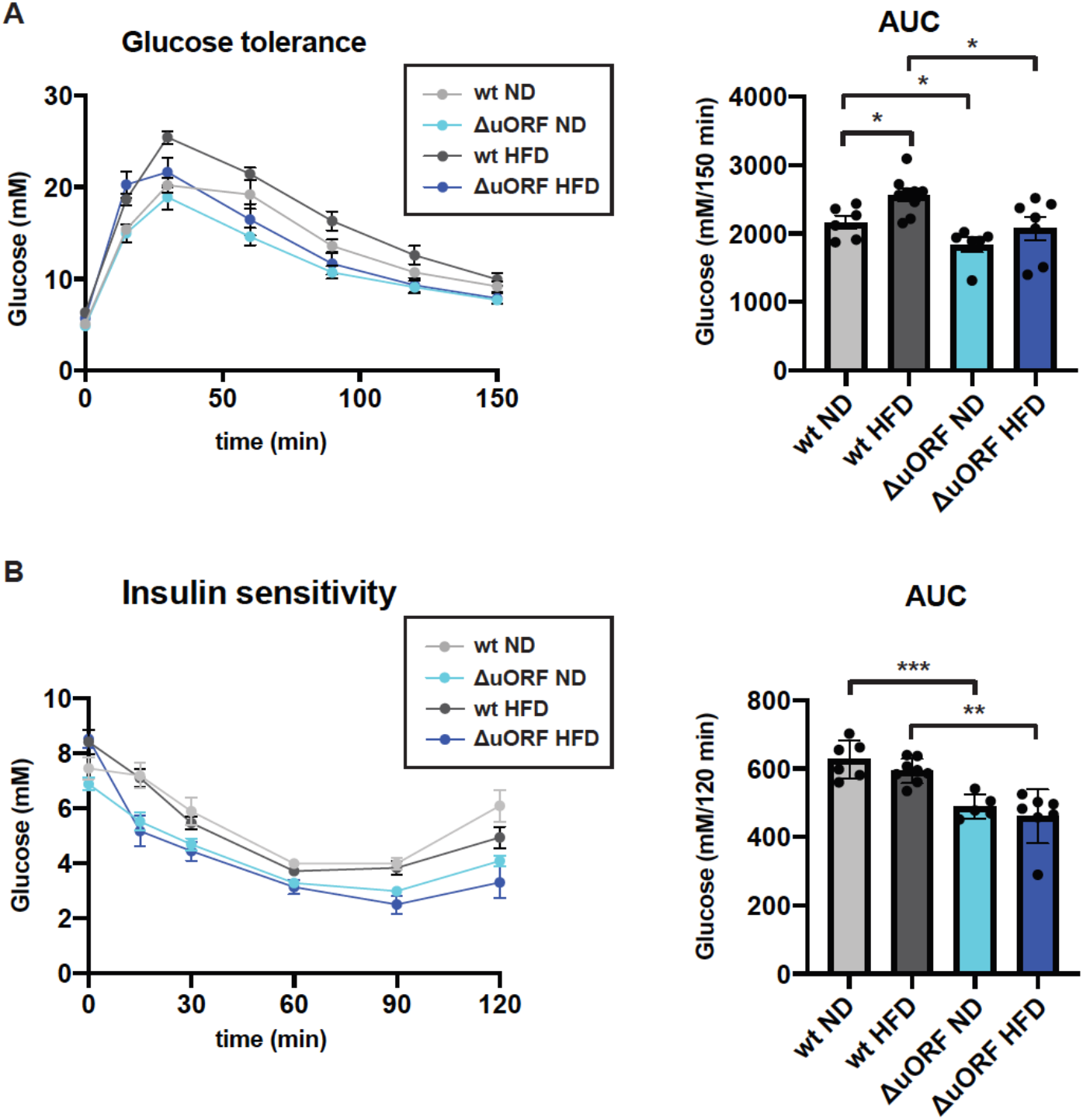
*Cebpb*^*ΔuORF*^ mice maintain glucose tolerance and insulin sensitivity on a high-fat diet (HFD). (A) Glucose tolerance test (IPGTT) with the calculated area under the curve (AUC) of *Cebpb*^*ΔuORF*^ mice (ΔuORF) and wt mice injected i.p. with glucose (2 g/kg) after a 16-h fast (7 weeks; wt, ND n = 6; HFD n = 9; ΔuORF ND n = 6; HFD n = 7). (B) Insulin sensitivity test (IPIST) with the calculated area under the curve (AUC) of fed of *Cebpb*^*ΔuORF*^ mice (ΔuORF) and wt mice injected i.p. with insulin (0.5 IU/kg) (6 weeks; wt, ND n = 6; wt, HFD n = 8; ΔuORF, ND n = 6; ΔuORF, HFD n = 7). All values are mean ± SEM. P-values were determined with Student’s t-test, *P < 0.05; **P <0.01; ***P < 0.001.

C/EBPβ is a known transcriptional regulator of fat cell differentiation and function (Siersbaek and Mandrup, 2011). We have shown earlier that the truncated C/EBPβ isoform LIP inhibits adipocyte differentiation and that fibroblasts derived from *Cebpb*^*ΔuORF*^ mice have an increased adipogenic differentiation potential. We therefore analyzed the expression of selected adipogenic factors in the visceral fat of by quantitative PCR. The expression of both key differentiation factors C/EBPα and PPARγ was clearly higher in *Cebpb*^*ΔuORF*^ mice on HFD compared to wt controls (**Figure 4**). In addition, the expression of sterol regulatory element-binding protein 1c (SREBP1c), a key factor for lipogenesis and fatty acid synthase (FAS), a key lipogenic enzyme, are higher in *Cebpb*^*ΔuORF*^ mice compared to wt (**Figure 4**). Taken together, these data suggest an improved fat storage function through loss of the inhibitory LIP isoform.

**Figure 4.**
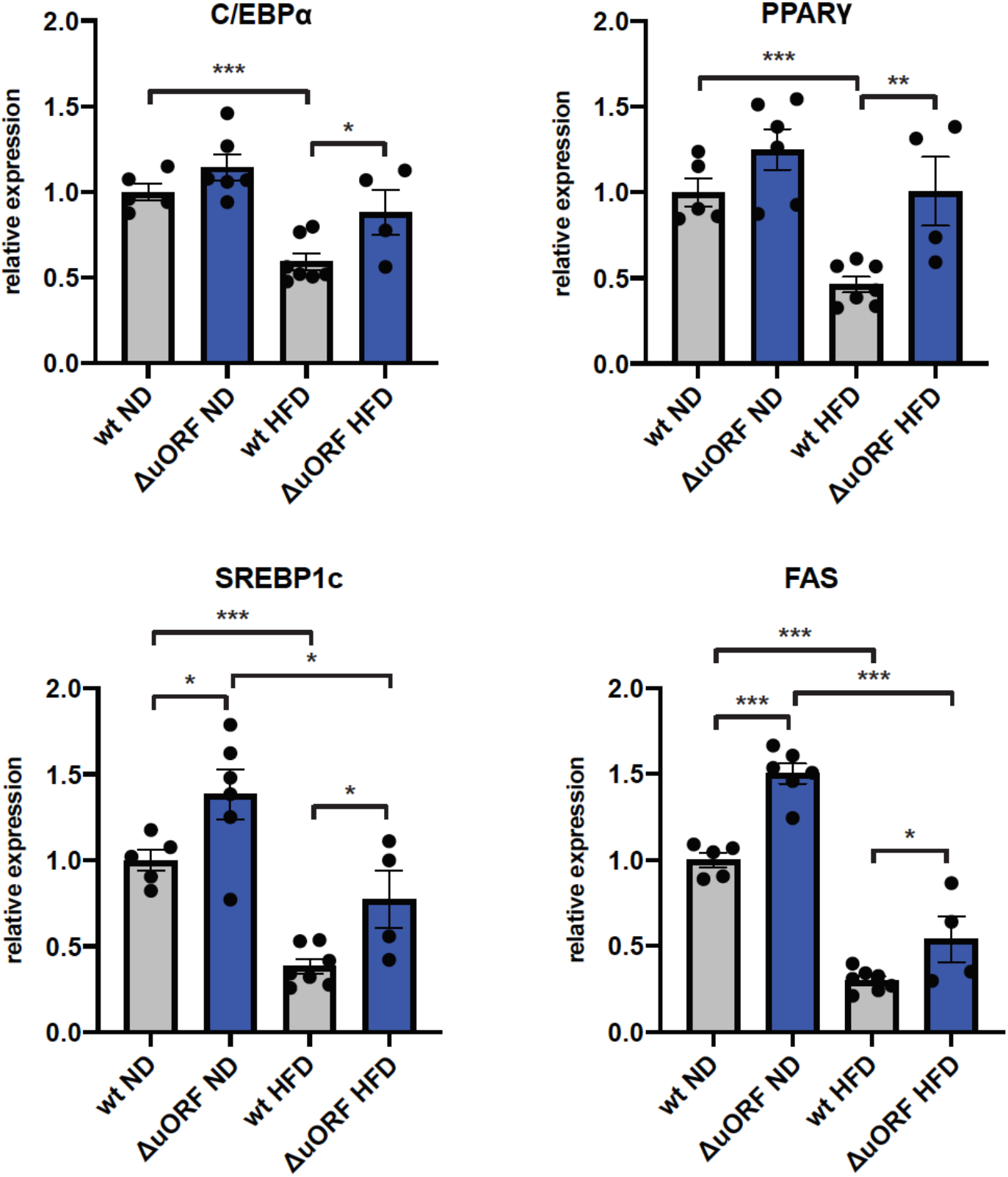
Expression of key adipogenic genes is elevated in *Cebpb*^*ΔuORF*^ mice on high-fat diet (HFD). Relative mRNA expression levels of the adipogenic transcriptionfactors C/EBPα, PPARγ and SREBBP1c and key enzyme FAS measured in livers of *Cebpb*^*ΔuORF*^ mice (ΔuORF) and wt mice on either normal diet (ND) or HFD (19 weeks; wt ND n = 5; wt HFD; n = 6; ΔuORF ND, n = 6; ΔuORF HFD, n = 4). All values are mean ± SEM. P-values were determined with Student’s t-test, *P < 0.05; **P < 0.01; ***P < 0.001.

## Discussion

In two previous reports we have shown that *Cebpb*^*ΔuORF*^ mice display metabolic improvements and a delay in the onset of age-related conditions, collectively resulting in an extended lifespan in females (Müller et al., 2018; Zidek et al., 2015). Here we demonstrate that *Cebpb*^*ΔuORF*^ mice are protected against the development of metabolic disturbances in response to HFD feeding. This improved metabolic phenotype occurs although the total fat mass in *Cebpb*^*ΔuORF*^ mice is more increased in response to HFD than in wt mice. We suggest that two special features of the white adipose tissue (WAT) in *Cebpb*^*ΔuORF*^ mice contribute to these metabolic improvements. Firstly, *Cebpb*^*ΔuORF*^ mice on a HFD store the surplus of nutrients in fat depots that expand through hyperplasia; they increase the number of adipocytes and thus the individual cells have to store less fat. These smaller adipocytes are metabolically more active and less inflamed compared to the inflated wt adipocytes residing in a hypertrophic fat depot. Hypertrophic adipocytes are known to secrete inflammatory cytokines that promote insulin resistance and other metabolic disturbances (Reilly and Saltiel, 2017; Weisberg et al., 2003). Furthermore, since the number of adipocytes in hypertrophic fat tissue does not increase and the amount of fat that can be stored in an adipocyte is limited, fat starts to accumulate in ectopic tissues like liver or muscle which compromises metabolic health (Frasca et al., 2017). Accordingly, we observed a pronounced dyslipidemia in wt mice on HFD but not in the *Cebpb*^*ΔuORF*^ mice. Secondly, the *Cebpb*^*ΔuORF*^ mice store relatively more fat in the subcutaneous compartment than wt mice, which relieves the fat storage pressure for the visceral depots. Furthermore, fat storage in the subcutaneous fat depot is associated with a better metabolic health status in humans and mice, while fat storage in the visceral fat depot is associated with insulin resistance and inflammation (Carey et al., 1997; McLaughlin et al., 2011; Tran et al., 2008).

The HFD induced adipocytic hyperplasia in *Cebpb*^*ΔuORF*^ mice indicates that unrestrained LAP functionality - through loss of inhibitory function of LIP - stimulates adipocyte differentiation and function. It may explain why *Cebpb*^*ΔuORF*^ mice store more fat in WAT on a HFD than wt littermates assuming that efficient fat storage by adaptive increase of the number of adipocytes prevents relocation of fat to peripheral tissues. These observations are in line with our previous experiments showing that mouse embryonic fibroblasts (MEFs) derived from *Cebpb*^*ΔuORF*^ mice are much more efficiently induced to undergo adipogenesis than wt MEFs, and differentiation of 3T3-L1 preadipocytes is strongly suppressed upon ectopic induction of LIP (see data in Expanded View Figure 3B, C of (Zidek et al., 2015)). Furthermore, the Kirkland lab has shown that overexpression of C/EBPβ-LIP in preadipocytes from young rats impaired adipogenesis, in context of a study showing that decline in C/EBP-function caused by and age-related decline in C/EBPα expression with concurrent upregulation of C/EBPβ-LIP affects adipogenesis (Karagiannides et al., 2001). Also, we and others have observed an age-associated increase in LIP expression (Hsieh et al., 1998; Müller et al., 2018; Timchenko et al., 2006).

In the current model of the regulatory cascade of adipocyte differentiation C/EBPβ and C/EBPδ induce the expression of C/EBPα and PPARγ, which by positive feedback stimulate each other’s expression (Siersbaek and Mandrup, 2011). Pharmacological activation of PPARγ by thiazolidines similarly to the *Cebpb*^*ΔuORF*^ mutation stimulates adipocyte differentiation, results in fat storage in hyperplastic adipocytes and a shift to fat storage in the subcutaneous compartment, resulting in improved metabolic health (Adams et al., 1997; Fujiwara et al., 1988; McLaughlin et al., 2010; Okuno et al., 1998). Our data propose pharmacological reduction of LIP expression as an additional approach to switch the unhealthy metabolic phenotype of obese individuals into a healthy obese phenotype to prevent the development of metabolic disease. We have shown that a search for such an intervention is feasible through the identification of drugs that affect LIP or LAP expression similar to mTORC1-inhibition (Zaini et al., 2017).

The metabolic disturbances induced by obesity superimposed on ageing significantly aggravates age-related metabolic decline and chronic inflammation (inflammaging) (Frasca et al., 2017). The aging resilience observed in *Cebpb*^*ΔuORF*^ mice associated with metabolic improvements together with the here demonstrated favorable metabolic response to HFD-feeding suggest that C/EBPβ is a central factor in the physiological response to nutritional/environmental changes and the malleability of the aging process. Various studies have revealed a crucial role of fat metabolism in mediating the positive effects of calorie restriction (CR) on health and lifespan. In particular, mice under CR were shown to burn more fat than they eat (Bruss et al., 2010). Ingested carbohydrates are first converted into fat via *de novo* lipogenesis in fat tissue, which subsequently is catabolized to generated fatty acids that are transported and used for energy production in peripheral tissues like liver and muscle. Our previous study suggest that C/EBPβ is involved in controlling this metabolic roundabout (Zidek et al., 2015). Furthermore, It was shown that maintaining an active fat tissue metabolism under CR is a prerequisite for lifespan extension in response to CR in mice (Liao et al., 2011), and that in Drosophila enhanced fatty acid metabolism is required for the positive effects of CR (Katewa et al., 2012). Taken together these studies and the recent discovery that prevention of C/EBPβ recruitment to its enhancers results in segmental progeria with impaired energy homeostasis and lipodystrophy (Schafer et al., 2018) demonstrate that C/EBPβ is an important factor in aging and lifespan determination (Niehrs and Calkhoven, 2020), and that fat metabolism is a crucial process involved.

## Material and methods

### Mice

*Cebpb*^*ΔuORF*^mice (Wethmar et al., 2010) were back-crossed for 6 generations into the C57BL/6J background. Mice were kept at a standard 12-h light/dark cycle at 22°C in a pathogen-free animal facility and for all experiments age-matched male mice were used. Mice were fed a High fat diet (HFD) (60% fat, D12492, Research Diets New Brunswick, USA) for 19 weeks starting at an age of 12 weeks or a standard chow diet (normal diet, ND) (10% fat, D12450B, Research Diets New Brunswick, USA) as control. For each genotype weight-matched mice were distributed over the different diet groups. Mice were analyzed at different time points as indicated in the figure legends. The determination of body weight and food intake (per cage divided through the number of mice in the cage) was performed weekly for 16 or 18 weeks, respectively. During the performance of all experiments the genotype of the mice was masked. All animal experiments were performed in compliance with protocols approved by the Institutional Animal Care and Use committee (IACUC) of the Thüringer Landesamt für Verbraucherschutz (#03-005/13).

### Determination of body composition

Mice were anaesthetized and the abdominal region from lumbar vertebrae 5 to 6 was analyzed using an Aloka LaTheta Laboratory Computed Tomograph LCT-100A (Zinsser Analytic) as described in (Zidek et al., 2015).

### Determination of caloric utilization

Both the feces and samples of the HFD food were collected, dried in a speed vacuum dryer at 60°C for 5h, grinded and pressed into tablets. The energy content of both the feces and food samples was determined through bomb calorimetry using an IKA-Calorimeter C5000. The energy efficiency was calculated through subtraction of the energy loss in the feces from the energy consumed.

### IP glucose tolerance and insulin sensitivity tests

For the determination of glucose tolerance, mice were starved overnight (16h) and a 20% (w/v) glucose solution was injected i.p., using 10 μl per gram body weight. After different time points the blood glucose concentration was measured using a glucometer (AccuCheck Aviva, Roche). For the determination of insulin sensitivity, an insulin solution (0,05 IU/ml insulin in 1xPBS supplemented with 0.08% fatty acid-free BSA) was i.p. injected into non-starved mice using 10μl per gram body weight and the blood glucose concentration was measured as described above.

### Histological staining

Tissue pieces were fixed for 24h with paraformaldehyde (4%) and embedded in paraffin. Sections (5μm) were stained with hematoxylin and eosin (H&E) in the Autostainer XL (Leica). For lipid staining with Sudan III, cryosections (10μm) were fixed with paraformaldehyde (4%) and stained with Sudan-III solution (3% (w/v) Sudan-III in 10% ethanol and 90% acetic acid) for 30 min.

### Determination of organ weight

After termination of the mice organs were collected and cleaned from surrounding fat or connective tissue and their weight was determined using an analytical balance.

### qRT-PCR analysis

Tissue pieces were homogenized using the Precellys 24 system (Peqlab) in the presence of 1ml QIAzol reagent (QUIAGEN). The RNA was isolated using the RNeasy Lipid Tissue Mini kit (QUIAGEN) according to the protocol of the manufacturer, incubated with RQ1 RNase-free DNase (Promega) for 30 min at 37°C and purified further using the RNeasy Plus Mini kit (QUIAGEN) starting from step 4.

1 μg RNA was reverse transcribed into cDNA with Oligo(d)T primers using the Transcriptor First Strand cDNA Synthesis kit (Roche). The qRT-PCR was performed with the LightCycler 480 SYBR Green I Master mix (Roche) using the following primer pairs: CD68: 5’-GCCCACCACCACCAGTCACG-3’ and 5’-GTGGTCCAGGGTGAGG GCCA-3’, PPARγ 5’-GCCCTTTGGTGACTTTATGG-3’ and 5’-CAGCAGGTTGTCTT GGATGT-3’, C/EBPα 5’-CAAGAACAGCAACGAGTACCG-3’ and 5’-GTCACTGGTC AACTCCAGCAC-3’, SREBP1c: 5’-AACGTCACTTCCAGCTAGAC-3’ and 5’-CCACT AAGGTGCCTACAGAGC-3’, FAS: 5’-ACACAGCAAGGTGCTGGAG-3’ and 5’-GTCC AGGCTGTGGTGACTCT-3’, and β-actin: 5’-AGAGGGAAATCGTGCGTGAC-3’ and 5’-CAATAGTGATGACCTGGCCGT-3’.

### Statistical methods

The number of biological replicates is indicated as n = x. All graphs show average ± standard error of the mean (SEM). To calculate statistical significance of the obtained results the Student’s t-Test was used with *p<0.05; **p<0.01 and ***p<0.001. Single mice were excluded when results indicated technical failure of the experimental performance. Furthermore, extreme outliers were excluded from the analysis.

## Acknowledgements

We thank Susanne Klaus and Susanne Keipert (DIfE, Potsdam) for help with bomb calorimetry. At the FLI, Verena Kliche for technical assistance, the staff of the animal house facility for embryo transfer and advice on mouse experiments, and Maik Baldauf for help with histology. L.M.Z. was supported by the Deutsche Forschungsgemeinschaft (DFG) through a grant (CA 283/1-1) to C.F.C.

